# AttentiveDist: Protein Inter-Residue Distance Prediction Using Deep Learning with Attention on Quadruple Multiple Sequence Alignments

**DOI:** 10.1101/2020.11.24.396770

**Authors:** Aashish Jain, Genki Terashi, Yuki Kagaya, Sai Raghavendra Maddhuri Venkata Subramaniya, Charles Christoffer, Daisuke Kihara

## Abstract

Protein 3D structure prediction has advanced significantly in recent years due to improving contact prediction accuracy. This improvement has been largely due to deep learning approaches that predict inter-residue contacts and, more recently, distances using multiple sequence alignments (MSAs). In this work we present AttentiveDist, a novel approach that uses different MSAs generated with different E-values in a single model to increase the co-evolutionary information provided to the model. To determine the importance of each MSA’s feature at the inter-residue level, we added an attention layer to the deep neural network. The model is trained in a multi-task fashion to also predict backbone and orientation angles further improving the inter-residue distance prediction. We show that AttentiveDist outperforms the top methods for contact prediction in the CASP13 structure prediction competition. To aid in structure modeling we also developed two new deep learning-based sidechain center distance and peptide-bond nitrogen-oxygen distance prediction models. Together these led to a 12% increase in TM-score from the best server method in CASP13 for structure prediction.

## Introduction

Computational protein structure prediction is one of the most important and difficult problems in bioinformatics and structural biology. Understanding protein structure can unlock information about protein function, and can aid in the design and development of artificial proteins and drug molecules^1,2^. Recently, a significant improvement in protein structure prediction has been observed due to improvements in contact and, more recently, distance map prediction^3^. The predicted contacts/distances are used to drive computational protein folding, where the 3D atomic protein structure is predicted without the need for template structures^4^.

The core principle behind modern contact prediction is detecting coevolutionary relationships between residues from multiple sequence alignments (MSAs)^5^. Previous contact map prediction approaches used direct coupling analysis to identify these relationships. These methods include CCMPred^6^, PSICOV^7^, Gremlin^8^, EV fold^9^, and plmDCA^10^. The next wave of methods, which represents the current state of the art, uses deep learning to predict contacts/distances. Deep learning-based methods have improved contact prediction significantly. This is evident from the recent community-wide assessment for structure prediction, CASP13^3^ (Critical Assessment of Structure Prediction), where top-performing methods in structure prediction including AlphaFold^11^ and methods in contact prediction including RaptorX^12^, TripletRes^13^, and ZHOU Contact^14^ are all deep learning-based. Raptor-X and Alphafold also showed that predicting distance distributions instead of binary contacts can further improve the performance. However, the current approaches are still not accurate enough to consistently achieve structure modeling with high GDT-TS structure evaluation scores^3^. Thus, further improvement is still needed.

One of the keys for accurate distance/contact prediction is the quality of MSAs^15,16^. Recent works have used a conservative E-value cutoff to generate MSAs because using a large E-value cutoff can lead to noisier and sometimes incorrect co-evolution information in the MSA. On the other hand, a larger E-value cutoff can yield an MSA containing more sequences, which may provide useful information particularly when a query protein does not have many close homologs. The difficulty is that the appropriate level of sequence similarity depends on the protein family^17,18^.

Here, we developed a new deep learning-based approach, AttentiveDist, which predicts protein distance maps with high accuracy. AttentiveDist uses a set of MSAs that are obtained with different E-value cutoffs, where the deep-learning model determines the importance of every MSA using an attention mechanism. Attention mechanisms in deep learning models are widely used in natural language processing^19,20^ and computer vision^21,22^ for determining which regions in the sentence or image respectively are important for a given task. To better generalize the model, we used a multi-tasking approach, predicting backbone angles and orientation angles^23^ together with inter-residue distance. We also show that structure prediction from a predicted distance map using Rosetta^24^ can be improved by using predicted inter-residue sidechain center distances and main-chain hydrogen-bonds. The predicted distances and angles are converted into potentials using neural network-predicted background distributions.

We show that AttentiveDist outperforms the best methods for contact prediction in CASP13. We compared distance predictions using combinations of individual MSAs of different E-value cutoffs with the attention-based approach, showing that the latter achieved a better precision. We also demonstrate that the attention given to different MSA-based features in AttentiveDist is correlated to the co-evolutionary information in the MSA. Finally, we show that AttentiveDist significantly outperforms the top CASP13 server models in structure prediction, with an average TM-score of 0.579 on 43 CASP targets compared to the 0.517 of the top server method in CASP13.

## Results

### AttentiveDist architecture

AttentiveDist predicts the distribution of Cβ - Cβ distance and three side-chain orientation angles for each amino acid residue pair, as well as backbone dihedral angles. Its uses a deep learning framework, ResNet^25^, with an attention mechanism that identifies important regions in MSAs.

**Figure 1** shows the network structure of AttentiveDist. The network is derived from ResNets^25^, where each residual block consists of convolution layers followed by instance normalization^26^ and exponential linear unit^27^ (ELU) as the activation function. This set is repeated twice with a dropout layer in between to form one residual block. The first 5 residual blocks are feature encoding layers and the weights are shared for the different inputs generated by 4 MSAs of E-values 0.001, 0.1, 1, and 10. For multiple different MSA feature encoding, we use soft attention to automatically determine the most relevant MSA for each pair of residues. An attention weight vector *a* of size *k* is computed for every *i, j* pair of residues, where *k* is the number of different MSAs used. Let *X_m_* be the encoded feature matrix for MSA *m. a_m_* is a scalar value that represents the “attention” or importance given to encoded feature *X_m_*(*_i,j_*), which is computed using Equation 1. The matrix W in Equation 1 is chosen such that *e_m_* is scalar, and it is learned during training along with the other parameters of the network. The attended feature matrix *Y* is computed as the weighted sum of different MSA encoded features where the weight is attention given as shown in Equation 2. The intuition is that Y captures the relevant information from multiple different MSAs.

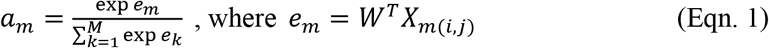

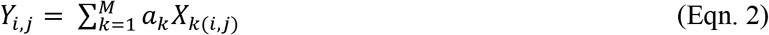

**Figure 1.**
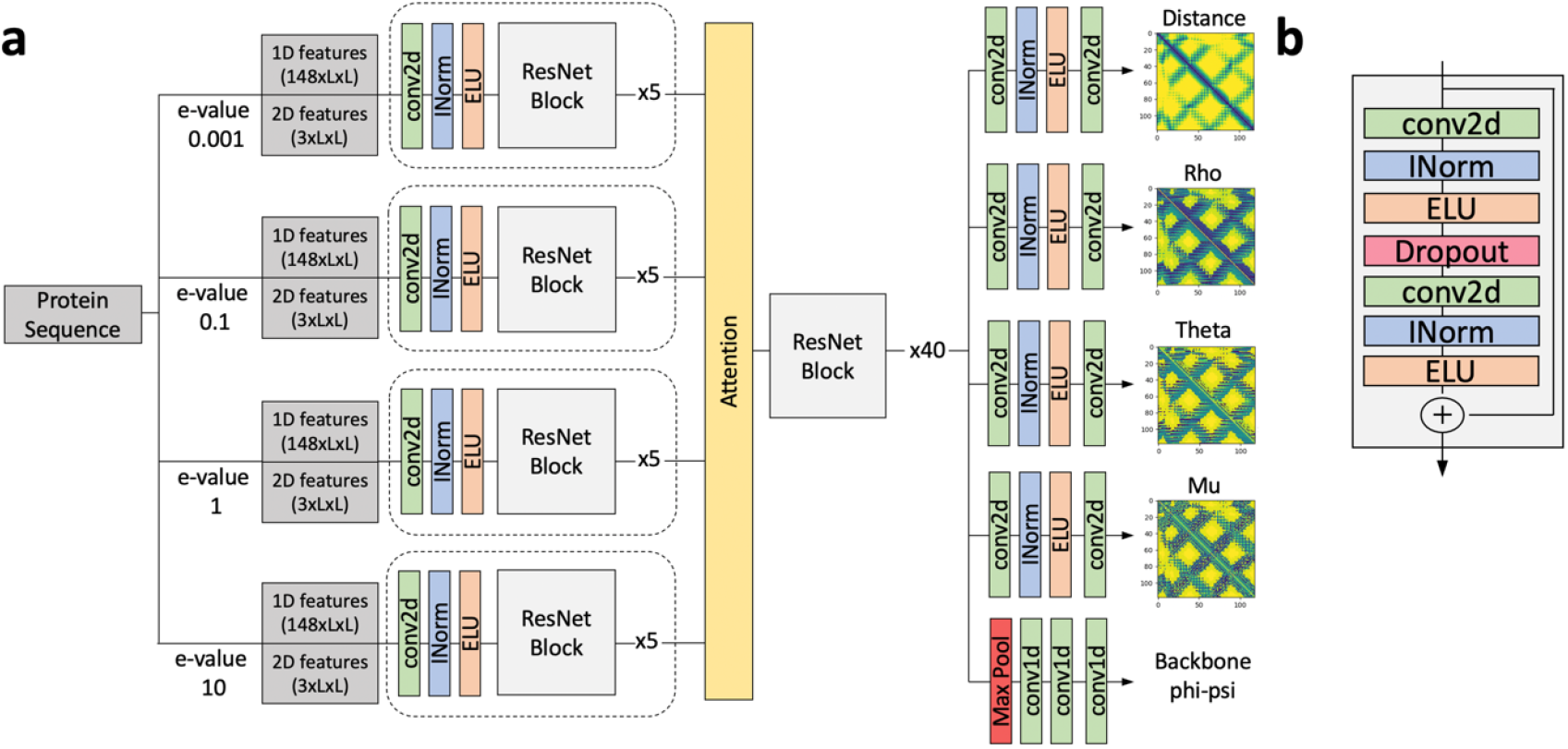
The network architecture of AttentiveDist. **a**. The overall architecture. From sequence-based features computed from a set of MSA’s of different E-values and 2D features, AttentiveDist uses ResNet with attention mechanism to predict Cβ-Cβ distances, three side-chain orientation angles, and backbone φ, ψ angles.. Dotted box represents weights are shared. **b**. Layers in a single ResNet Block. conv2d (green), 2d convolution layer; INorm (blue), instance normalization; ELU (orange), Exponential Linear Unit.

The attended features are then passed through 40 residual blocks. The model branches into 5 different paths with different outputs after the last residual block. In each path there is an additional convolution layer followed by normalization and activation which learn task-specific representations. To improve the generalization, we used a multi-task learning approach where the model is trained on six related tasks, namely, distance prediction, three side-chain orientation angles (**Supplementary Figure 1**), and the φ, ψ backbone angles. The paths for distance and orientation angles contain a final convolution layer to obtain the proper output dimension, followed by softmax activation. In the backbone φ, ψ angles path, a max pooling layer is added to reduce the dimensionality from LxLx64 to Lx64 where L is the size of the protein, followed by 1D convolution and softmax activation. The whole network is trained end-to-end. The final model is an ensemble of 5 models, where the prediction is the average of individual E-value models and the attention-based model that combines the four MSAs.

We used eight sequence-based input features. The 1D features are one hot encoding of amino acid type (20 features), PSI-BLAST^28^ position specific scoring matrix (20 features), HMM^29^ profile (30 features), SPOT-1D^30^ predicted secondary structure (3 features) and solvent accessible surface area (1 feature), making a total of 74 1D features. MSAs, from which the 1D features were computed, were generated using the DeepMSA^15^ pipeline. 1D features were converted into 2D features by combining features of two residues into one feature vector. We also used three 2D features, which were a predicted contact map by CCMPRED^6^ (1 feature), mutual information (1 feature), and statistical pairwise contact potential^31^ (1 feature). Thus, in total we used (2 x 74) + 3 = 151 L x L features, where L is the length of the protein.

The AttentiveDist network predicts the Cβ – Cβ distance of every pair of residues in a target protein as a vector of probabilities assigned to 20 distance bins. The first bin is for 0 to 4 Å, the next bins up to 8Å are of a size 0.5 Å and then bins of a 1 Å size follow up to 18 Å. The last bin added is for no-contact, i. e. for 18 Å to an infinite distance. Similarly, the backbone φ, ψ angles were binned to 36 ranges, each of which has a 10-degree range. Three side-chain orientation angles, ρ, θ, and μ (**Supplementary Figure 1**) were binned into 24, 24, and 15 bins, each with a size of 15 degrees, respectively. The side-chain orientation angles were only considered between residue pairs that are closer than 20 Å, and for the rest of the residue pairs a no contact bin was considered as the correct answer. For target values for training, the real distances and angles were converted into vectors where the bin containing the real distance/angle has value 1 and while the rest were set to 0.

The network was trained on a dataset of 11,181 non-redundant proteins, which were selected from the PISCES^32^ database. Sequences released after 1st May 2018 (i.e. the month of beginning of CASP13) were removed. Every pair has less than 25% sequence identity. Out of these, 1,000 proteins were selected randomly as the validation set, while the rest were used to train the models. More details are provided in the Method section.

### Contact prediction performance on the CASP13 Dataset

We compared the performance of AttentiveDist with the top-performing methods in CASP13 on 43 FM (Free Modeling) and FM/TBM (Template-Based Modeling) domains. FM and FM/TBM are harder targets compared to template-based modeling because they do not have any appropriate template protein available, necessitating de-novo prediction. We used the standard metric of top L/n predicted long range contacts precision as used in other works, where L is length of the protein and n is 1, 2, and 5. Long range contacts are defined as contacts between residues that are 24 or more residues away. Since AttentiveDist predicts residue-residue distances instead of binary contact, we converted this to contact prediction by summing the probabilities of distance bins from minimum distance to 8 Å.

**Table 1** summarizes the results. The first 3 rows show the performance of top methods in CASP13, RaptorX-Contact^12^, TripletRes^13^, and ZHOU Contact^14^, all of which are deep learning-based approaches. AttentiveDist outperforms the best CASP13 methods in L/1, L/2 and L/5 precision. There was a significant improvement in L/1 precision of 2.5% when compared to Raptor-X. Most of the deep learning-based methods are ensembles of multiple methods. AttentiveDist uses 5 models whereas other methods use much higher numbers of models. For example, Raptor-X is an ensemble (average) of 28 models. We further show that for most of the domains, AttentiveDist improves the performance of L/1 precision over Raptor-X, the best among the three existing methods compared (**Supplementary Figure 2**). AttentiveDist showed a higher L/1 precision than Raptor-X for 23 domains, and tied for 2 domains out of the 43 domains.

**Table 1.** CASP13 FM and FM/TBM 43 targets long range precision.

| **Model** | **L/5** | **L/2** | **L/1** |
| --- | --- | --- | --- |
| ZHOU Contact | 0.662 | 0.543 | 0.426 |
| TripletRes | 0.701 | 0.587 | 0.451 |
| RaptorX-Contact | 0.744 | 0.612 | 0.481 |
| AttentiveDist | **0.746** | **0.624** | **0.493** |

L/5, L/2 and L/1 shows precision values when top L/5, L/2 or L/1 contact predictions with the highest probabilities were considered where L is the length of the protein.

### Model Ablation Study

We performed an ablation study of our model to understand how much different additions contribute to the performance (**Table 2**). The baseline model shown at the top of the table is a single model that predicts only Cβ-Cβ distance using an E-value of 0.001 for feature generation. 0.001 was used for E-value because it gave the overall the highest precision among the other E-values used in AttentiveDist. Next, we added multitask learning, where the model predicts the distance, 2D side-chain orientation angles, and the backbone dihedral angles together, but without attention. The multi-task learning improved the L/1 precision from 0.451 to 0.468.

**Table 2.** Ablation study on the 43 CASP13 FM and FM/TBM domain targets

| Model | L/5 | L/2 | L/1 |
| --- | --- | --- | --- |
| Distance only (E: 10<sup>-3</sup>) | 0.700 | 0.586 | 0.451 |
| E-value 10<sup>-3</sup> | 0.716 | 0.608 | 0.468 |
| E-value 10<sup>-1</sup> | 0.693 | 0.587 | 0.452 |
| E-value 1 | 0.724 | 0.589 | 0.455 |
| E-value 10 | 0.696 | 0.580 | 0.452 |
| No-attention, 4 E-values | 0.705 | 0.602 | 0.472 |
| Attention, 4 E-values | 0.716 | 0.613 | 0.479 |
| AttentiveDist (Ensemble) | 0.746 | 0.624 | 0.493 |

The next three rows compare multi-task learning results with four different E-values (0.001, 0.1, 1, and 10). The results show that on average an E-value of 0.001 performed the best. The sixth row, “No attention, 4 E-values” shows the results of using four E-values to compute four different 1D features but without the attention mechanism. This increased the L/1 precision to 0.472. When the attention mechanism was further introduced, the L/1 precision, further increased it from 0.472 to 0.479. Finally, we averaged the outputs from the 5 models, which is AttentiveDist that resulted in 0.14 gain to achieve 0.493 in L/1 precision.

### Prediction performance relative to the size of MSAs

As observed by previous works^12,13^, we also observed correlation between the size of MSAs, i.e. the number of sequences in the MSAs and the contact prediction accuracy. In **Figure 2a,** the L/1 long range contact precisions were shown for two methods, AttentiveDist and the model using only MSAs of E-value 0.001, relative to the number of sequences in the MSAs. The number of sequences in the MSAs is shown in the log scale. A positive correlation was observed, as shown in the figure, and particularly, there is clear distinction of the performance at the sequence count of 100. When the sequence count was less than 100, L/1 precision was always below 0.4. Oppositely, when the sequence count is very high, over 10,000, high precisions of over 0.75 were observed. Although the high precision was observed with a large number of sequences, observed precisions had a large range of values when the sequence counts was over 100.

**Figure 2.**
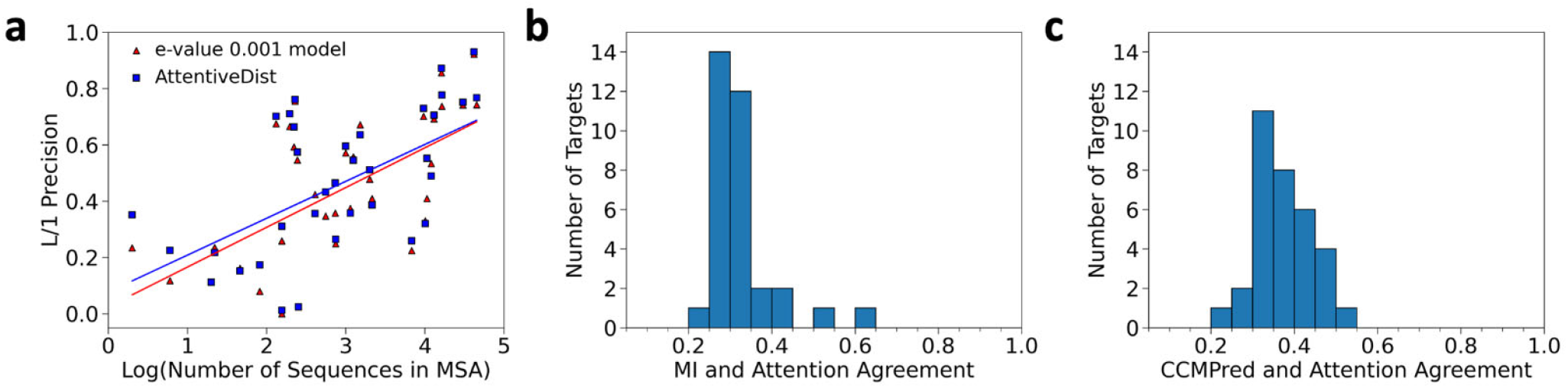
Analysis of the MSA size and the attention. **a,** Relationship between log of the sequence counts in MSAs and long-range L/1 contact precision for the 43 CASP13 targets. AttentionDist (blue) and the E-value 0.001 model (red), where E-value 0.001 was used as a cutoff for generating MSAs. The lines represent the regression. **b**, the fraction of residue pairs where the MSA with the highest attention agreed with the MSA with the highest mutual information (MI). The number of targets among the 35 CASP13 target proteins that have the particular fraction of agreed residue pairs were counted for each bin. 43 FM and FM/TBM CASP13 target domains belong to 35 proteins. Out of the 35 proteins, two proteins were discarded from this analysis because the four MSAs with different E-value cutoffs of these proteins were identical. **c**, the agreement is compared with the contact probability computed from the four MSAs with CCMPred.

### Analyses of attention weights

In AttentiveDist, for each residue pair, attention values are distributed across four MSA-based features each computed with the four different E-value cutoffs, which sum up to 1.0. To understand what the attention mechanism captures, in **Figure 2b** and **2c** we examined how the attention corresponds to co-evolution signals. We compared with local and global co-evolutionary signals. The local co-evolutionary signal used is mutual information (MI), which uses pairwise residue profile information. The global signal considers effects from other residues as well, which can be computed by pseudo-likelihood maximization (PLM). We used CCMPred^6^, which is an implementation of PLM. For each residue pair in a protein target, we counted the number of times the MSA with the highest attention weight assigned by AttentiveDist agrees with the MSA with the highest co-evolutionary signal.

We observed that for both MI and CCMPred the agreement was higher than random (0.25). The histogram shifted to higher values when compared with CCMPred than MI. The average agreement for the 33 proteins were higher for CCMPred (0.376) than MI (0.329). Thus, overall the attention is a mechanism to select MSAs with higher co-evolutionary signals.

### Protein structure modeling

Finally, we built the tertiary structure models of the CASP13 domains and compared with the top CASP13 server models. For the structure modeling, in addition to the predictions of Cβ-Cβ distance, main-chain φ, ψ angles, and the three, ρ, θ, and μ, side-chain orientation angles, we tested the inclusion of two additional distance constraints, which were Side-chain CEnter (SCE)-SCE distances and peptide-bond nitrogen (N)-oxygen (O) atom distances. These distances help in proper secondary structure formation and side-chain packing. All the constraints were converted into a potential function by normalizing predicted probability values in bins by predicted reference probability values. The folding was performed using Rosetta^24^ by adding the predicted potentials into the Rosetta energy function. Out of a few thousand models generated, the best scoring model for each target are reported in this section. Details are provided in Methods.

The average TM scores of our approaches and the top-three CASP13 severs, Zhang-Server, RaptorX-DeepModeller, and BAKER-ROSETTASERVER are shown in **Figure 3a**. For AttentiveDist, we showed results by two versions, one with the predicted SCE-SCE distances and the backbone N-O distances, which is denoted as AttentiveDist (Full), and the one without these two distance constraints (AttentiveDist w/o SCE and N-O). AttentiveDist (Full) outperformed the other methods, achieving an average TM score of 0.579 while the top CASP13 servers, Zhang-Server, RaptorX-DeepModeller, and BAKER-ROSETTASERVER had values of 0.517, 0.508, and 0.450, respectively. Comparing two versions of AttentiveDist, the two distance constraints improved the TMscore by 2.1% from 0.568. In **Supplementary Figure 3**, TM-scores of individual domain targets by the two versions of AttentiveDist are compared. In **Figure 3b**, we further show the TMscores of the 43 individual targets by AttentiveDist (Full) and Zhang-Server. AttentiveDist (Full) showed a higher TM-Score than Zhang-Server for 29 cases and tied for 3 cases.

**Figure 3.**
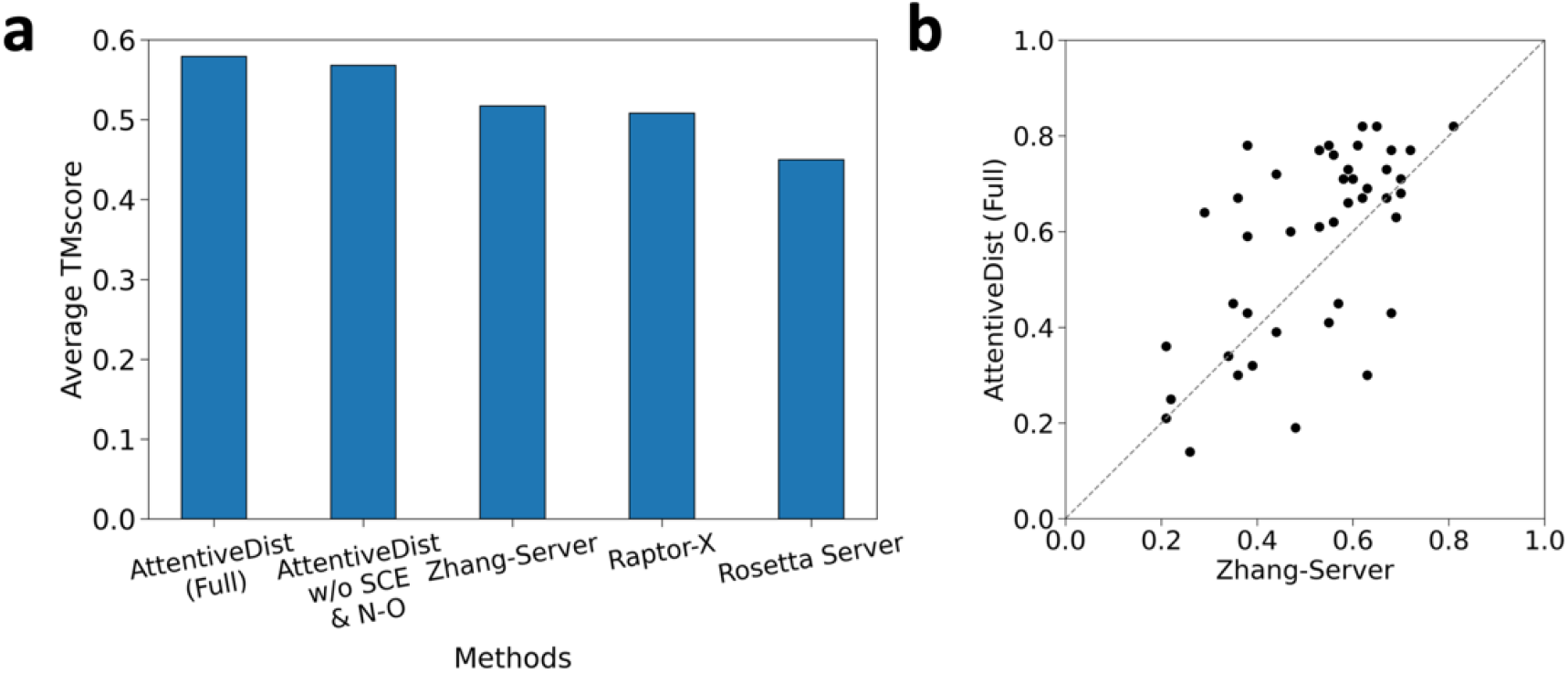
Performance in structure modelling. **a**, TM-score for AttentiveDist, AttentiveDist without using predicted sidechain center distance and backbone N-O distance and the top 3 server methods in CASP13 for 43 FM and FM/TBM targets. **b**, Individual target TM-score comparison between our method and the Zhang-Server. The registered name of Raptor-X in CASP13 was RaptorX-DeepModeller and BAKER-ROSETTASERVER for Rosetta Server.

**Figure 4** provides four examples of models computed with distance prediction by AttentiveDist (Full) in comparison with Zhang-Server, RaptorX-DeepModeller, and BAKER-ROSETTASERVER. The first panel, **Figure 4a**, is a 180-residue long domain with two α-helices and two β-sheets, T0957s1-D1. While our model has a TM-score of 0.78, indicating that the overall conformation is almost correct, the models by the other three methods have some substantial differences from the native. The Zhang-Server model missed one β-sheet, the RaptorX-DeepModeller did not predict any β-sheets, and the BAKER-ROSETTASERVER placed the β-sheet at the top of the structure and a α-helix in substantially different orientations. The second example, T0980s1-D1 (**Figure 4b)** is another αβ class protein with a long loop region, which is placed on the right-hand side of the figures. The loop is difficult to correctly model, as the three top CASP13 servers did not fold it well. The incorrect modeling of the loop also affected to the placement of the α-helix in the right orientation in their models. Our AttentiveDist model managed to have the overall fold almost correct, as shown by a higher TM-score of 0.64. For the next target, T0986s2-D1 (**Figure 4c**), the Zhang-Server has almost all the architecture correct, but slight shifts of α helices cost it in the TM-score, which was 0.59. Our model had the conformation almost correct even in the loop regions, resulting in a high score of 0.78. The BAKER-ROSETTASERVER model did not assemble the large β-sheet correctly. The last target shown has an α-helical structure, which consists of two long α-helices with multiple small α-helices. (T0950-D1, **Figure 4d**). While our model identified correct orientations for the two long helices, the other methods built them incorrectly which caused other incorrect helix arrangements at the top of the structure in the figure, resulting in lower scores.

**Figure 4.**
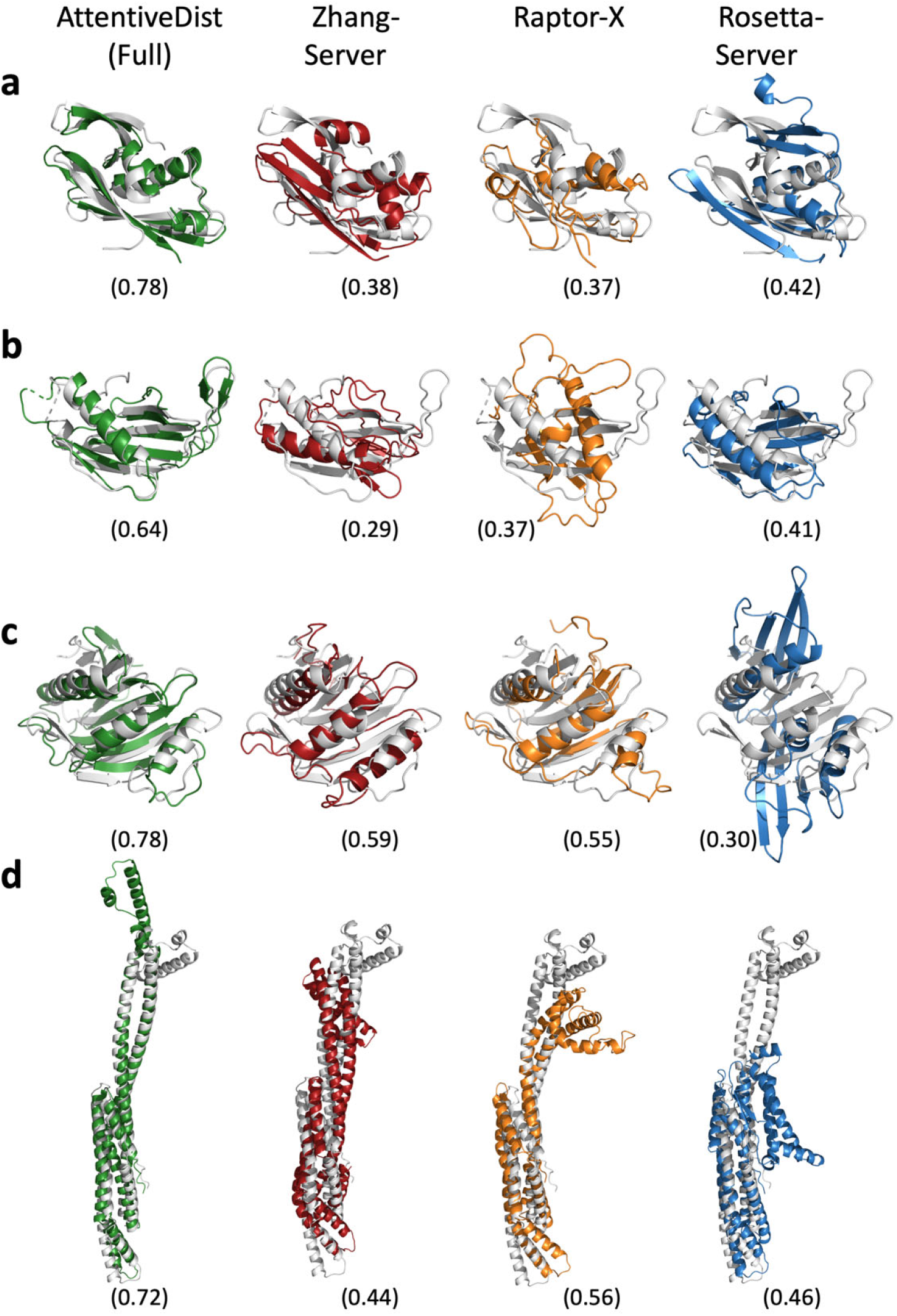
Examples of structure models by AttentiveDist (Full) in comparison with the top-1 model by the three top servers. AttentiveDist (Full), green; Zhang-Server, red; RaptorX-DeepModeller, orange; and BAKER-ROSETTASERVER, blue. The native structures are shown in gray. TM-scores are shown in parentheses. Targets are **a,** T0957s1-D1 (PDB ID: 6cp8; length: 180 amino acids); **b**, T0980s1-D1 (PDB ID: 6gnx; 104 aa); **c**, T0986s2-D1 (PDB ID: 6d7y; 155 aa); **d**, T0950-D1 (PDB ID: 6ek4; 331 aa).

## Discussion

We presented AttentiveDist, a deep learning-based method for predicting residue distances and angles from four MSAs with four different E-value cutoffs. By adding an attention layer to the network, useful features from MSAs were selectively extracted, which led to competitive performance relative to the top performing server methods in CASP13. In AttentiveDist, the attention layer as well as multi-tasking strategy boosted the prediction accuracy. In the context of the recent intensive efforts for developing residue distance/contact prediction methods by the community, this work shows another strong demonstration of how protein structure information can be further squeezed by exploiting modern deep learning technologies. Although our approach showed improvement over existing methods, an improvement is still needed especially when the available sequences are sparse for input MSAs, which remains as an important future work.

## Methods

### Training, validation, and test datasets

For the training and validation dataset, we took proteins from the PISCES^32^ database that consists of a subset of proteins having less than 25% sequence identity and a minimum resolution of 2.5 angstroms, released in October 2019. We further pruned this dataset by removing proteins that contain more than 600 or less than 50 amino acids and those released after 1st May 2018 (i.e. the month of beginning of CASP13). Next, proteins that have intermediate gaps of more than 50 residues, not considering the termini, were removed. Finally, a protein that has 2-letter chain names was removed because PISCES capitalizes chain names making it confusing for cases where the real 2 letter chain name has both mixed lowercase and uppercase alphabets used. This resulted in 11,181 proteins. Out of those, 1,000 proteins were selected randomly as the validation set, and the rest were used to train the models. For each instance of glycine, a pseudo-Cβ atom was built to be able to define Cβ-Cβ distance by converting it to alanine.

CASP13 FM and FM/TBM domains were used as the test set, containing 43 domains (across 35 proteins). The full protein sequence was used in the input instead of the domain to replicate the CASP13 competition.

### MSA generation

To generate the MSA we used the DeepMSA^15^ pipeline. This pipeline consists of three stages where three different databases are searched to obtain similar sequences, which produces better MSAs compared to a single database search. The packages used for DeepMSA were HHsuite^33^ version 3.2.0 and HMMER^34^ version 3.3. The sequence databases we used were released before the CASP13 competition began for the sake of fair comparison, and were: Uniclust30^35^ database dated October 2017, Uniref90^36^ dated April 2018, and Metaclust_NR^37^ database dated January 2018. We generated 4 different MSAs with E-value 0.001, 0.1, 1, and 10 used in HHsuite^33^ and HMMER^21,34^.

### Network parameters and training

In AttentiveDist the convolution filter (kernel) size is 5×5 for the first 3 blocks and then 3×3 for the rest of the network, and the channels were kept constant to 64. We also added dilation to increase the receptive field of the network, with dilation cycling through 1,2 and 4.

The loss function used during training is the weighted combination of individual objective loss. For each objective cross-entropy loss was used and the weights were manually tuned. Distance and orientation angles losses were given weight of 1 while the backbone φ and ψ angle losses were given weight of 0.05 each. The Adam^38^ optimizer with a learning rate of 0.0001 was used. Dropout probability was 0.1. Dilations were cycled between 1,2 and 4. The learning rate, dropout and loss weights were tuned on the validation dataset. We trained the model for 80 epochs. Batch size was set to 1 because of GPU memory constraints.

### Sidechain center distance and backbone hydrogen-bond (N-O) prediction

For the tertiary structure modeling, we tested the inclusion of two additional predicted distance constraints, distances between Side-Chain cEnters (SCE) and distances between the nitrogen and the oxygen (N-O) in peptide bonds. These distances were binned similarly to the Cβ – Cβ distances. The first bin was for a distance between 0 to 2 Å, bins up to 20 Å were of a width of 0.5 Å, followed by a bin of size 20 Å to infinite. A bin for residue pairs with missing information was also added. For prediction, we used networks with 25 ResNet blocks, which is smaller than the one in **Figure 1.** The model was trained on the E-value 0.001 MSA data (**Supplementary Figure 4**). The prediction performance is show in **Supplementary Table 1.**

### Protein 3D structure generation from distance prediction

We performed protein structure modeling similar to the work by Yang et al. ^23^ We used Rosetta’s protein folding and energy minimization protocols with customized constraints. The constraints were computed from our predictions of distance distributions (Cβ-Cβ, SCE-SCE, and backbone N-O) and angle distributions (backbone φ–ψ and the three residue-pair orientation angles) by normalizing the predicted values with predicted reference distributions. For both distance and angle constraints, the predicted distributions were converted to an energy potential as follows:

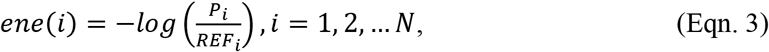

where *P_i_* and *REF_i_* are the predicted probability and the reference probability of *i*-th bin, respectively. *N* is the number of bins in the predicted distribution.

The reference probability distributions of three distances, backbone angles, and the side-chain orientation angles were predicted with a five-layer fully-connected neural networks. A network of the same architecture was trained for each type of constraints. For a distance type, the features used were the positions *i* and *j* of the two amino acids, the length of the protein, and a binary feature of whether a residue is glycine or not^11^. For angle predict we also included the one-hot encoding of the amino acid type.

All energy potentials were smoothed by the spline function in Rosetta, and then used as constraints in the energy minimization protocol. The energy potentials of distances (Cβ-Cβ, SCE-SCE and backbone N-O) and inter-residue orientations were split into L/10*L/10 blocks. To explore a diverse conformational space, the blocks of the potentials were randomly added to the energy function in the minimization steps. We generated 4,000 decoy models with different folding paths (i.e. additions of the blocks of potentials) and weight parameters that balance the energy terms. All decoy models were ranked by ranksum^39^, a sum of the ranks of three scoring functions, GOAP^40^, DFire^41^, and ITScore^42^. The best scoring model was selected as the predicted structure.

## Acknowledgments

We thank Xiao Wang for checking and test running the codes. This work was partly supported by the National Institutes of Health (R01GM123055, R01GM133840) and the National Science Foundation (DMS1614777, CMMI1825941, MCB1925643, DBI2003635).

## Author Contributions

D.K. conceived the study. A.J. developed the algorithm and wrote the codes. A.J., Y.K. and S.R.M.V.S. generated MSAs. G.T. generated protein structures models. C.C. computed score ranking of generated protein structure models. A.J., G.T., and D.K. analyzed the results. A.J. drafted and D.K. edited and finalized the manuscript.

---

**Supplementary Figure 1.**
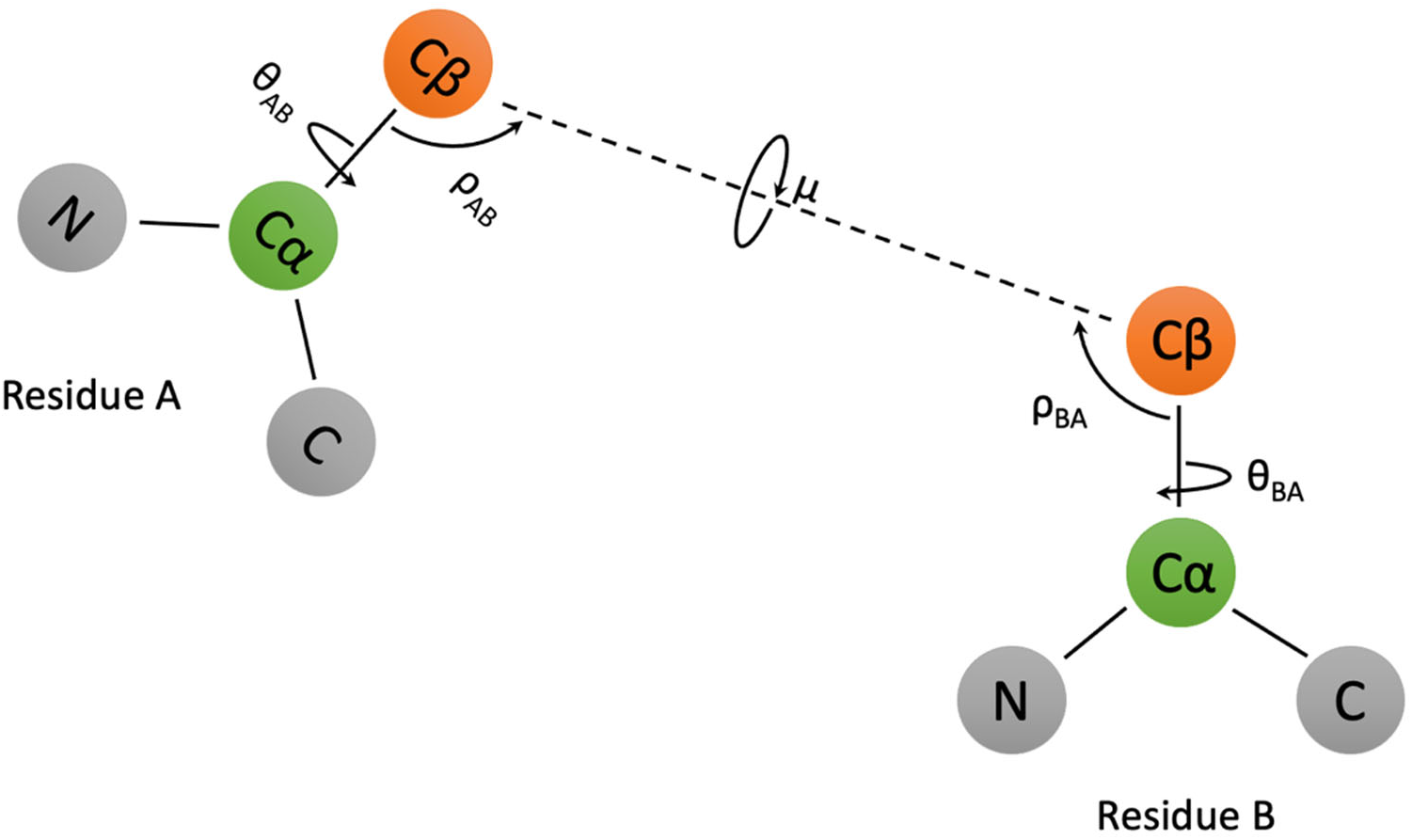
Orientation angles. The three orientation angles mu(μ), theta(θ) and rho(ρ) between any pair of residues in a protein. In the 3d structure of the protein, considering any two residues A and B, θ_AB_ represents the dihedral angle between the vectors N_A_ -> Cα_A_ and Cβ_A_ -> Cβ_B_ along the axis of Cα_A_ -> Cβ_A_. ρ_AB_ represents the angle between the vectors Cβ_A_ -> Cα_A_ and Cβ_A_ -> Cβ_B_. θ and ρ depends on the order of residue and thus are asymmetric. μ represents the dihedral angle between the vectors Cα_A_ -> Cβ_A_ and Cβ_A_ -> Cα_B_ along the axis of Cβ_A_ -> Cβ_B_. These orientation angles help in representing the direction of residue A to residue B and vice-versa. The orientation angles were originally described in Yang et al ^1^. We replaced the symbols from the paper by Yang et al. to prevent confusion with conventionally used angle notation.

**Supplementary Figure 2.**
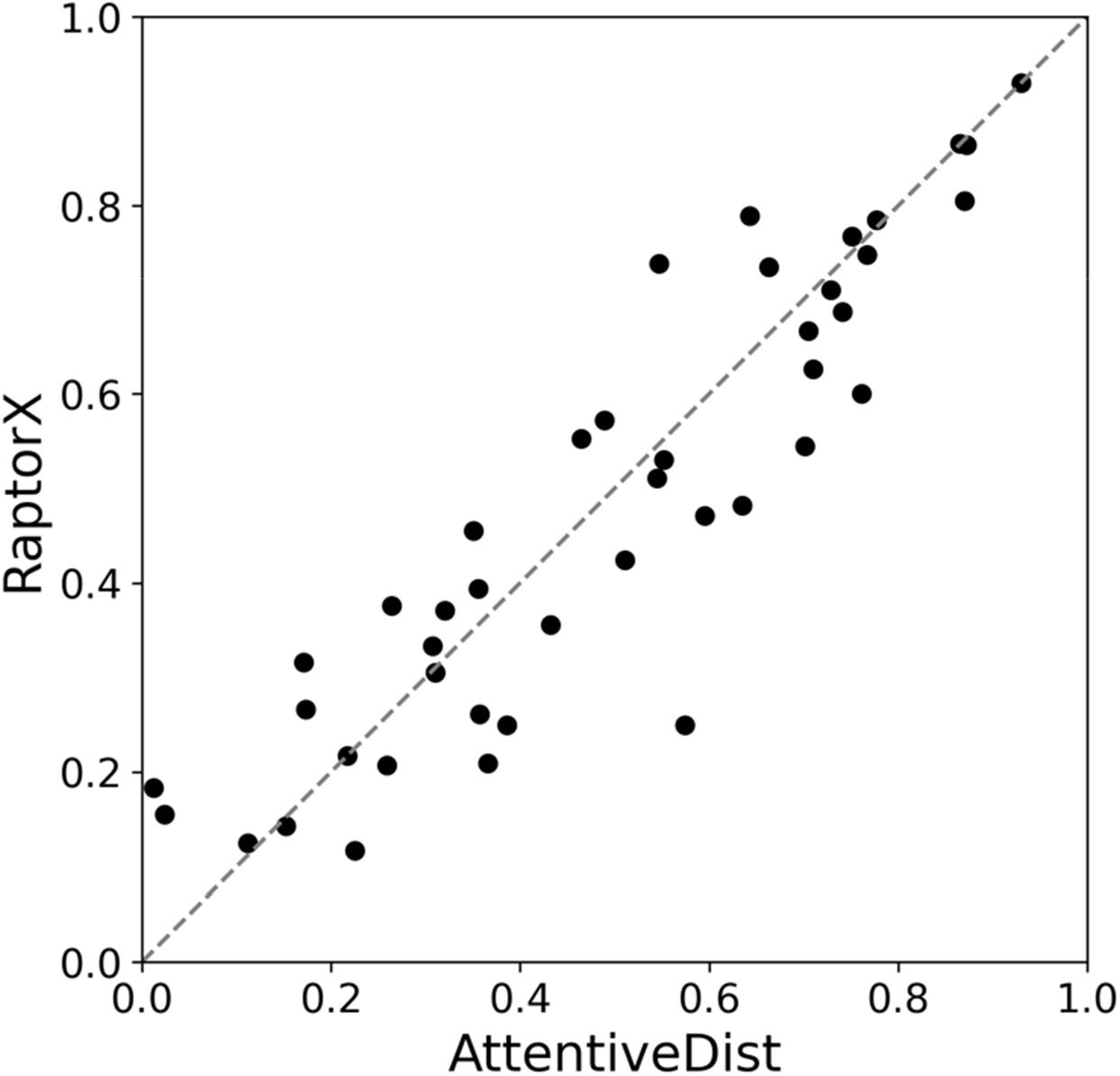
Long L/1 precision comparison between Raptor-X and AttentiveDist on the CASP13 FM and FM/TBM domains. AttentiveDist showed a higher L/1 precision than Raptor-X for 23 domains and tied for 2 domains out of the 43 domains.

**Supplementary Figure 3.**
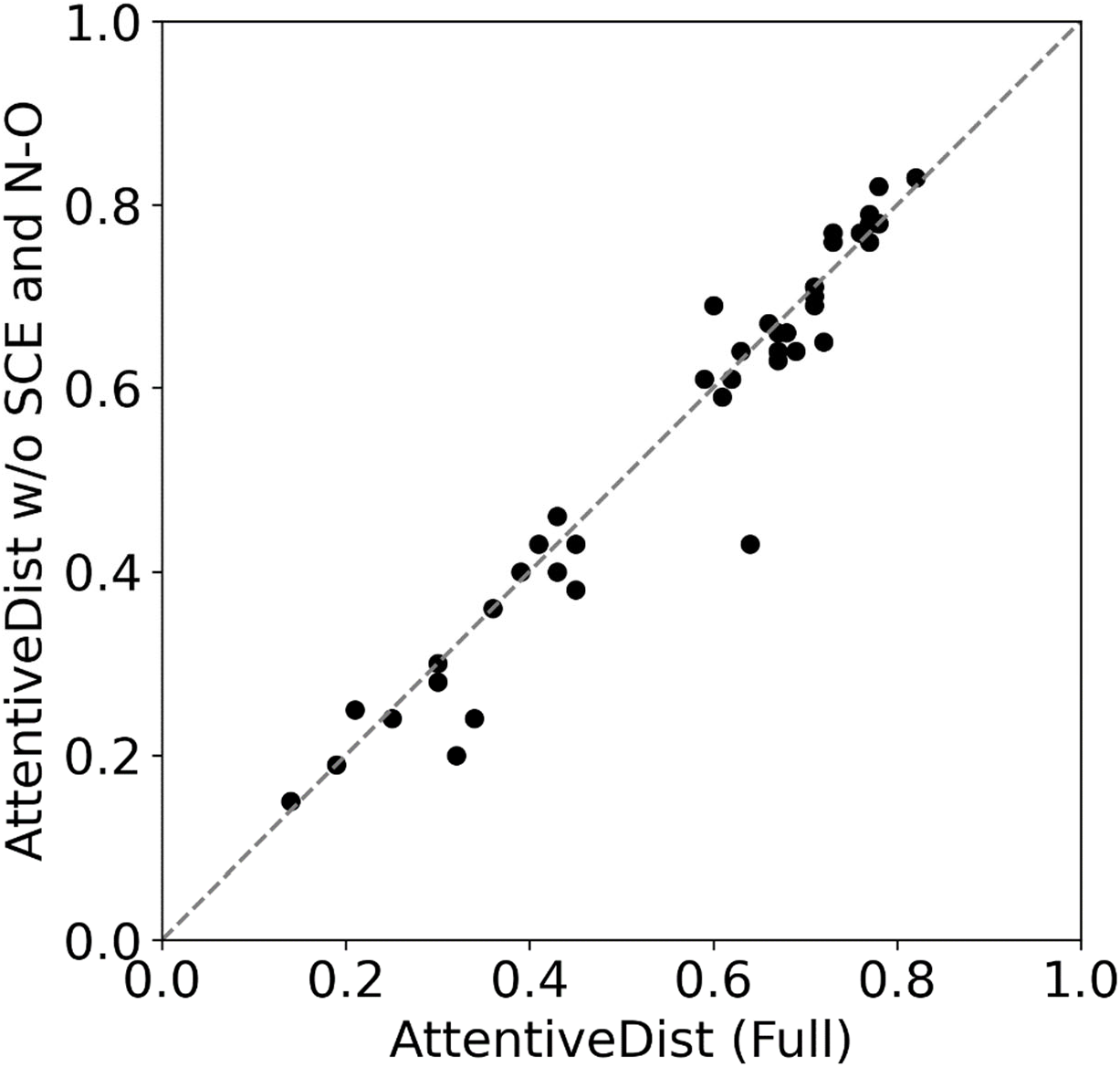
TM-score of AttentiveDist (Full) and AttentiveDist without using predicted SCE-SCE and N-O distances on the 43 CASP13 domains. AttentiveDist (Full) showed higher TM-Score for 19 targets, tied on 6 targets.

**Supplementary Figure 4.**
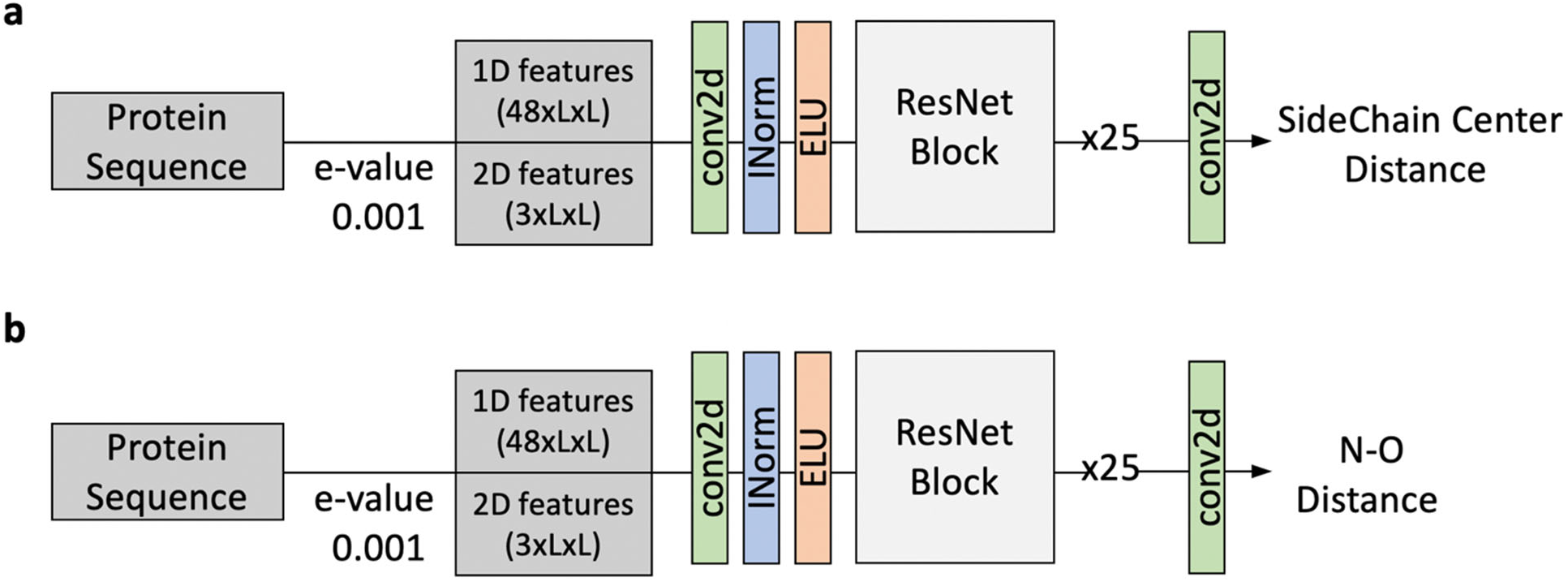
ResNet model architecture for **a**, Sidechain Center (SCE) distances and **b**, backbone peptide N-O pairwise distance prediction. The ResNet Block is the same as described in Figure 1b. conv2d (green) is 2d convolution layer, INorm (blue) is instance normalization, ELU (orange) is Exponential Linear Unit.

**Supplementary Table 1.** Accuracy of backbone phi-psi and orientation angles for the 43 CASP13 FM and FM/TBM domain targets. The bin size of torsional angles was set to 10° while the bin for the orientation angles was 15°. Bin slack of 0 represents that the predicted bin of the highest probability and the real bin were the same. Bin slack of 1(or 2) denotes that the predicted bin was 1(or 2) bin(s) away from the correct bin.

| Angle | Bin Slack |  |  |
| --- | --- | --- | --- |
| | 0 | 1 ($\pm 10^\circ$) | 2 ($\pm 20^\circ$) |
| $\varphi$ | 0.277 | 0.590 | 0.722 |
| $\phi$ | 0.242 | 0.548 | 0.700 |
| | 0 | 1 ($\pm 15^\circ$) | 2 ($\pm 30^\circ$) |
| $\mu$ | 0.332 | 0.582 | 0.634 |
| $\theta$ | 0.377 | 0.656 | 0.703 |
| $\rho$ | 0.394 | 0.692 | 0.748 |

**Supplementary Table 2.**
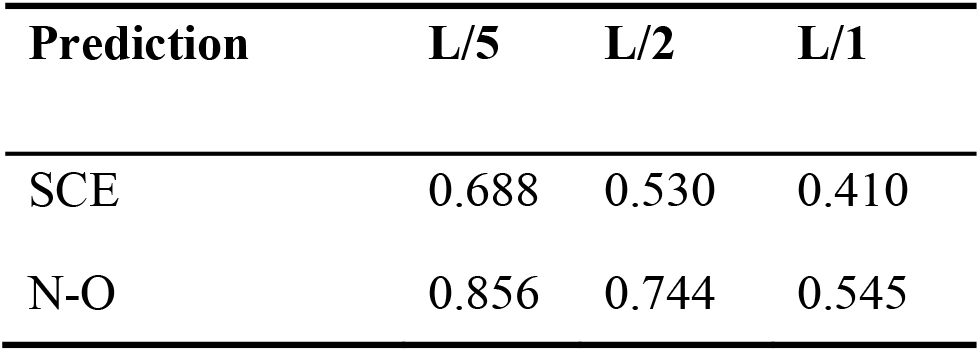
Long range precision of prediction made for Side-Chain cEnters (SCE) contact and contact between the nitrogen and the oxygen (N-O) in peptide bonds. The contact is defined if as pairs within 8 Å for SCE-SCE, and 4 Å for N-O. The 43 CASP13 FM and FM/TBM targets were considered.

